# Long-term effects of early-life adversity on DNA methylation in zebra finches

**DOI:** 10.1101/2025.04.23.649997

**Authors:** Marianthi Tangili, Blanca Jimeno, Michael Briga, Per J. Palsbøll, Simon Verhulst

## Abstract

Early-life experiences can have profound and long-lasting effects on adult phenotype and thereby Darwinian fitness, though the mechanisms driving these effects remain poorly understood. Epigenetic alterations, especially DNA methylation which affects gene expression, potentially mediate developmental condition effects on adult phenotype. We tested for such effects using captive zebra finches that were reared in either small or large broods, a manipulation that is known to have pleiotropic phenotypic effects. Analyzing whole genome DNA methylation patterns in erythrocytes from 50 individuals sampled in adulthood, we found 0.8% of all CpG sites after filtering to be differentially methylated after correction for multiple testing. We identified 149 non-transiently differentially methylated sites (DMSs) where the DNA methylation difference between treatments was larger than 25%. These DMSs were located in 19 autosomal chromosomes, in or near genes involved in critical biological processes such as cell growth, division, and differentiation, regulation of immune response, muscle contraction, and neuronal signaling. These findings suggest that epigenetic modifications such as DNA methylation potentially mediate long-term effects of early-life adversity via differential gene expression, but follow-up studies are needed to identify the extent to which the observed DMSs are functionally related to the previously observed phenotypic effects.

## Introduction

Early-life experiences can have long-lasting effects on adult phenotype, a phenomenon that is observed across the tree of life. In humans, early-life adversity has been associated with an increased life-long risk of developing psychopathological disorders and chronic health problems (McLaughlin, 2016). Similarly, in non-human primates, early-life adversity has been found to impact social relationships, survival, physical and mental health (Conti et al., 2012; Lange et al., 2023). In birds, early-life adversity can lead to altered mass regulation and foraging behavior (Andrews et al., 2015), attractiveness to mates (Holveck & Riebel, 2010; Mainwaring et al., 2012), social behavior (Gerritsma et al., n.d.) as well as accelerated cellular senescence and inflammation (Nettle et al., 2017).

The term phenotypic plasticity is used to describe all types of environmentally induced phenotypic variation (Stearns, 1989). Developmental phenotypic plasticity can occur prenatally through parental effects (e.g. maternal hormones, Schwabl, 1993), and postnatally through the conditions experienced by the individual during development (e.g. postnatal exposure to adversity, Spencer et al., 2009). However, the molecular mechanisms mediating the effects of early-life experiences on adult phenotype remain elusive. Environmentally-induced changes in the epigenome have emerged as important mechanisms for embedding early-life experiences in the genome and facilitating long-term effects (Aristizabal et al., 2020; Ruuskanen, 2024; Szyf, 2013; Szyf & Bick, 2012) leading to phenotypic plasticity (Hu & Barrett, 2017; Zhang et al., 2013). The most widely studied epigenetic modification is DNA methylation (DNAm), the addition of methyl groups to, typically, cytosine residues (CpG sites) throughout the genome. DNAm is a dynamic process that occurs during development and throughout the lifespan, playing a key role in regulating gene expression (Moore et al., 2013) while being significantly influenced by the environment (Law & Holland, 2019).

Advances in next generation sequencing and epigenetic methodologies have facilitated the exploration of the effect of early-life experiences on DNAm in multiple species and tissue types (Watamura & Roth, 2019). Early-life effects on the epigenome have been studied extensively in humans (Vinkers et al., 2015), showing that adverse, early-life experiences can leave long-lasting effects on DNAm. Early-life effects on the epigenome can in turn alter cellular/neuronal plasticity (Labonte et al., 2012), immune and developmental regulation (Bush et al., 2018) and even shape transgenerational biological embedding of adverse experiences (Scorza et al., 2023). In rats, maternal separation was associated with DNAm changes (Anier et al., 2014). Differential rearing led to altered DNAm patterns in the brain and T cells (Provençal et al., 2012). A recent review summarized the effects of multiple early-life biotic and abiotic environmental factors on avian epigenetic markers (Ruuskanen, 2024), pointing to the need to verify the causal links between epigenetic changes and phenotypic traits.

In altricial species, which depend on parental provisioning for early-life sustenance, an artificially enlarged brood size is an ecologically relevant approach to induce early-life stress. Being raised in a large brood increases sibling competition (Godfray, 1995), leading to increased begging intensity (Tangili et al., 2025) and altering parental provisioning rates (Bowers et al., 2014). Increased competition for resources within the nest is also known to affect growth (Nilsson & Svensson, 2001; Tangili et al., 2025), immunocompetence (Naguib et al., 2004; Saino et al., 2003), energy metabolism (Mertens, 1969) and, ultimately, lifespan (Briga et al., 2017). Previous research has shown that being reared in an enlarged brood influences the epigenetic regulation of genes related to growth, development, metabolism, behavior and cognition in great tit (*Parus major*) nestlings (Sepers et al., 2021). Being reared in an enlarged brood positively correlated with overall methylation levels in nestling (Sheldon et al., 2018) and enhanced the DNAm of the glucocorticoid receptor gene in adult zebra finches (*Taeniopygia guttata*, Jimeno et al., 2019). Yet, the long-term effects of an experimentally increased brood size of rearing on avian genome-wide DNAm levels remains unexplored. We employed whole genome enzymatic methyl-sequencing (EM-seq), to identify CpG sites where the level of DNAm in erythrocytes differed significantly among adult zebra finches raised in either small or large broods. If DNAm mediates the effects of early-life stress on adult phenotype, the level of DNAm in or near genes that contribute to phenotypic variation should differ between birds raised in small and large broods.

## Methods

### Brood size manipulation

Zebra finch pairs were randomly matched and placed in breeding cages (L x H x D: 80 × 40 × 40 cm) on a 14 L : 10 D schedule at ∼ 25^°^C temperature and ∼60% humidity. In the cages, there were nestboxes, nesting material (hay), food and water *ad libitum*. Fortified canary food (Bogena, Hedel, the Netherlands) was supplied to the parents until the hatching of the first chick in the nest to aid with reproduction. Nestboxes were checked daily for the presence of eggs or chicks. Chicks were cross-fostered until the maximum age of five days to broods of either two (*n*=25) or six (*n*=25) chicks, which is the range of brood sizes observed in the wild (Zann, 1996). On the day of cross-fostering, all chicks were weighed and individually marked by taking out some tufts located on their head. Siblings were always assigned to different non-parental broods, i.e., no parents were care takers of their own offspring to rule out genetic and parental phenotypic quality effects.

### Blood samples

Blood samples were collected the birds coming from manipulated broods during adulthood after they were translocated to outdoor aviaries (320 × 150 × 210 cm), each containing single sex groups between 18–24 adults (Briga et al., 2017). One hundred blood samples were collected between 2007 – 2015 from 50 known-age individuals (26 males and 24 females) sampled twice during their lifetime. The average sampling interval was 993 days. Blood samples were stored in glycerol storage buffer (40% glycerol, 50mM TRIS, 5mM MgCl2, 0.1mM EDTA) at -80^°^C.

### Enzymatic Methyl-seq

We extracted DNA according to the manufacturer’s protocol using innuPREP™ DNA Mini Kit (Analytik Jena GmBH) from 3 uL of nucleated red blood cells. Next-generation sequencing was conducted by the Hospital for Sick Children (Toronto, Canada). DNA was quantified using a Qubit High Sensitivity Assay. Two hundred ng of DNA was used as input material for library preparation using the Next Enzymatic Methyl-seq™ Kit (New England Biolabs Inc.) following the manufacturer’s instructions. Briefly, DNA was fragmented by sonication to an average length of ∼500bp using a Covaris LE220 (Covaris Inc.). Fragmented DNA was end-repaired and adapters ligated. 5-methylcytosines and 5-hydroxymethylcytosines were oxidized by TET2 and cytosines were deaminated by APOBEC. Methyl-seq libraries were subjected to five PCR amplification cycles. Libraries assessed on a Bioanalyzer™ DNA High Sensitivity chip (Agilent Inc.). The amount of DNA was quantified by quantitative PCR using the Kapa Library Quantification Illumina/ABI Prism™ kit according to the manufacturer’s instructions (La Roche Ltd.). Libraries were pooled in equimolar quantities and sequenced to a minimum of ∼100M paired-end reads (2×150bp) per sample on a 10B flow cell using a NovaSeqX™ (Illumina Inc.) platform.

### Bioinformatic processing of Enzymatic Methyl-seq data

Sequences were trimmed using Trim Galore! v0.6.10 (Krueger et al., 2023) in paired-end mode. The data were assessed before and after trimming using FastQC v. 011.9 (Andrews, 2010) and MultiQC v. 11.14 (Ewels et al., 2016). Alignments were performed using Bismark v. 0.14.433 (Krueger & Andrews, 2011) using the Bowtie 2 v. 2.4.5 alignment algorithm (Langmead & Salzberg, 2012) for both *in silico* bisulfite conversion of the reference genome and alignments. Trimmed reads were aligned against the *in silico* bisulfite converted zebra finch (*Taeniopygia guttata*) reference genome (GCA_003957565.4, Rhie et al., 2021). The average mapping efficiency was at 65.14% (SD: 2.79).

### Differentially methylated site (DMS) identification

Differentially methylated sites (DMSs) were identified using *Methylkit* v. 1.18.0 (Akalin et al., 2015). Since each individual was sampled twice, we merged the two observations per individual by summing their numbers of Cs, Ts and coverage using a custom R script. We calculated methylation as (Number of Cs) / (Number of Cs + Number of Ts) per individual. We filtered for a minimum coverage of 12 and maximum coverage of 99.9 to avoid PCR bias (Wreczycka et al., 2017). To normalize read coverage distributions between samples we used the *normalizeCoverage* function of the *Methylkit* package. The *unite* function was employed to retain CpG sites present in ≥70% of the sampled individuals. This threshold also implied the exclusion of the W chromosome. We calculated DNAm percentage/individual/site and excluded CpG sites at 0% or 100% DNAm in all samples (*n*=12,072). We calculated DNAm difference per CpG site using the *calculateDiffMeth* function using Fisher’s exact test (Fisher, 1922) and Bonferroni correction (Dunn, 1961). Following Wreczycka et al., (2017) we defined DMSs as CpG sites where the difference in percent methylation between individuals in the two treatments was at least 25%. With employing a very conservative approach to identify DMSs, we suspect that the likelihood of false positives has been greatly reduced.

### DMS annotation

The GFT-formatted annotation files for the zebra finch (GCF_003957565.2) were converted into BED12 format using the University of California, Santa Cruz (UCSC) utilities *gtfToGenePred* and *genePredToBed* (available at https://hgdownload.soe.ucsc.edu/downloads.html#utilities_downloads). CpG sites retained after filtering and DMS were annotated using the tool *annotateWithGeneParts* in the package *genomation* v. 1.4.1 (Akalin et al., 2015). This tool hierarchically classifies the sites into pre-defined functional regions, i.e., promoter, exon, intron, or intergenic, hereon referred to as annotation categories. The predefined functional regions were based on the annotation information present in the BED12 files accessed with the *genomation* tool *readTranscriptFeatures*. Annotations were performed as a hierarchical assignment (promoter > exon> intron> intergenic). Subsequently, a customized R script was employed to integrate each CpG site with their respective annotation category information.

We assessed the observed and expected proportion of DMS per annotation category using *χ*^2^ tests. The proportion of expected DMS per annotation category was estimated as the proportion of all CpG sites captured by our analysis located in each annotation category. For all *χ*^2^ tests, we applied a Benjamini-Hochberg correction to control for the false discovery rate to account for multiple hypotheses testing (Benjamini & Hochberg, 1995) and used an exact binomial test as a *post hoc* test to assess observed vs. expected occurrences. All statistical tests were carried out in R v. 4.1.2 (R Core Team, 2023).

### Gene ontology analysis

We ran a gene ontology analysis of 83 unique genes in whose promoters, exons and introns our identified DMS were located. For this we used the *ClueGo* plug-in v. 2.5.10 (Bindea et al., 2009) for *Cytoscape* v. 3.10.2 (Shannon et al., 2003) in order to identify gene ontologies (GO) that were significantly enriched. We selected zebra finch annotations along with the cellular component, molecular function, biological process ontologies and Kyoto encyclopedia of genes and genomes (KEGG, Kanehisa et al., 2002). For our analysis, we required GO terms to include a minimum of three associated genes. We applied a two-sided enrichment-depletion test using the Benjamini-Hochberg correction for multiple testing (Benjamini & Hochberg, 1995). GO categories were considered significantly enriched if their corrected p-value was less than 0.05. We opted to include all identified DMSs, regardless of genomic region. Although methylation in promoter regions is most commonly associated with gene silencing, DNAm in other genome regions can potentially affect gene expression as well through diverse mechanisms.

## Results

### Differentially methylated sites (DMSs)

The CpG-site-specific methylation difference between individuals from the two treatment groups was identified to be persistent throughout adulthood. This persistence was evidenced by comparable CpG-site-specific methylation differences between the treatment groups in both young and old individuals (Fig.S1). Taking that into consideration, we decided to proceed with the pooled reads from the two samples collected from the same individual to identify DMSs.

The CpG-site-specific DNAm difference between individuals raised in small and large broods ranged from -36.22 to 33.99% (Fig.1). After Bonferroni correction, 16,529 (0.8%) of the 2,052,610 sites retained after initial filtering (see Methods) were found to exhibit significant differential methylation between individuals raised in small and large broods (points above the horizontal dashed line in Fig.1). A total of 149 of the filtered sites (Table S1) fulfilled the criterion of a ≥|25%| methylation difference between individuals raised in small and large broods (Fig.2), and these were therefore classified as the differentially methylated sites (DMSs, blue points in Fig.1). The 149 DMSs were distributed among 19 autosomal chromosomes (Fig.2), with chromosome 29 showing the highest proportion of DMS that were significantly differentially methylated (Fig.S2), but note that this concerns only one DMS on a small chromosome. Chromosome 3, the third largest chromosome, harbored the second highest proportion of sites identified as DMS and the highest absolute number of DMS (*n*=102, Fig. S2). Coverage levels of CpG sites on chromosome 3 were not exceptionally high or low (Fig. S3,4), and the elevated proportion of DMS on chromosome 3 can therefore not be attributed to a higher (or lower) sequencing depth. Instead, we propose that this pattern may point to the biological importance of the genes located on chromosome 3.

**Figure 1.**
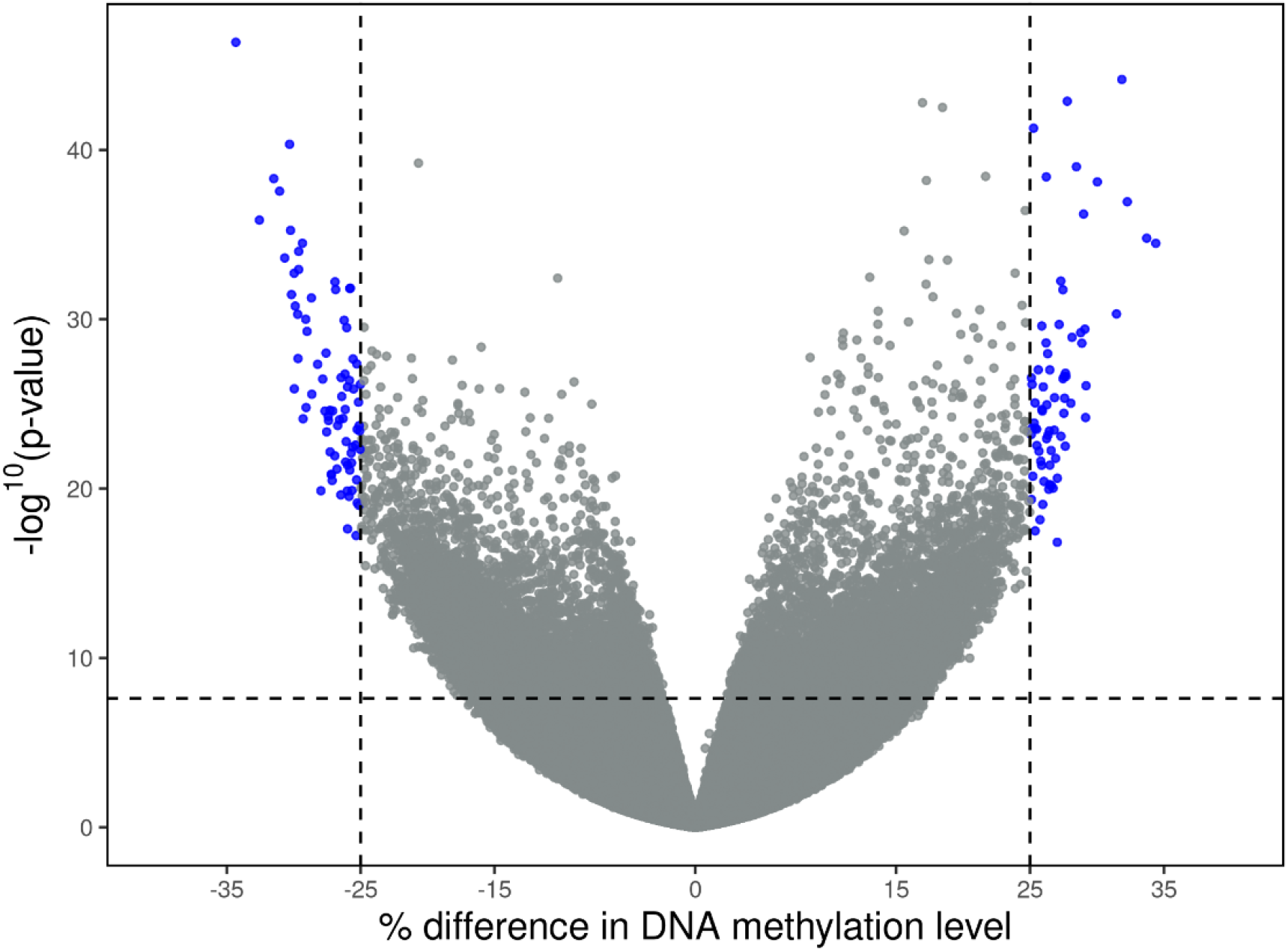
Volcano plot of Bonferroni-corrected p-values as a function of % DNAm differences between individuals raised in small and large broods. Each point represents a CpG site, with blue indicating biologically and statistically significant differences in % DNAm between individuals raised in small and large broods. The dashed horizontal line marks the genome wide significance threshold while the dashed vertical lines represent absolute 25% difference in DNAm.

Overall, the level of DNAm among the identified DMSs was not systematically higher or lower with respect to natal brood size. DNAm was higher in zebra finches raised in large broods at 79 DMSs, and lower in 70 DMSs. A more quantitative comparison yielded the same result: when we split the DMSs in sites with increased DNAm in individuals raised in large broods (median ± SE values, large: 56.00 ± 0.55; small: 30.63± 0.53,) and those with increased DNAm in individuals raised in small broods (small: 64.30 ± 0.48; large: 33.33 ± 0.52), these differences canceled each other out. Thus, DNAm differs between birds reared in small or large broods at many sites, but methylation averaged over all DMSs did not differ noticeably (mean ± SE, small =46.4±0.4, large =44.5±0.4, Fig. S5).

### Location of DMS in annotation categories and genes

Only one of the 149 identified DMSs was located in a promoter region, 83 DMSs were located in introns, 22 in exons and 43 in intergenic regions (Fig. 3). Exact binomial tests revealed that the proportion of DMSs observed was significantly lower and higher than expected by chance in intergenic regions (*p*=0.04) and introns (*p*=0.04), respectively. The observed and expected proportion of DMSs located in exons and promoter regions did not significantly differ from random expectations (introns: *p*=0.5; promoters: *p*=0.2).

The 149 CpG sites with methylation difference between brood sizes of ≥|25%| and a significant Bonferroni-corrected *p*-value, we located in or near 114 unique genes, of which 31 were uncharacterized (Table S1). Natal brood size was found to be associated with epigenetic changes affecting diverse biological, molecular and cellular processes. We detected ten enriched Gene Ontology (GO) terms (FDR-corrected *p*-value<0.05). The identified enriched GO terms were related to cell growth, division, and differentiation, regulation of inflammatory response, muscle contraction and neuronal function (Table S2).

## Discussion

Long-term effects of early-life conditions on adult phenotype and Darwinian fitness are well-documented, but the mechanisms mediating such effects remain largely unknown. DNAm influences gene expression and it can be modulated by the environment, potentially mediating the long-term effects of early-life conditions on life-long fitness. Support for this hypothesis comes from a wide range of taxa. In humans, CpG sites whose DNAm was affected by early-life conditions were associated with later-in life health conditions (Li et al., 2022). In spotted hyenas, early-life maternal care affected DNAm and stress physiology during later life stages (Laubach et al., 2021). In wild baboons, early-life resource limitation was found to be associated with DNAm patterns in adulthood (Anderson et al., 2024). These studies were all correlational, but experimentally manipulated early-life conditions have also been shown to affect DNAm in nestling zebra finches and great tits (Sepers et al., 2021; Sheldon et al., 2018). Jimeno et al. (2019) is, to our best knowledge, the only avian study that confirmed experimentally that DNAm effects of early-life conditions persist in adulthood, with cascading phenotypic effects, but their analysis was limited to a single gene (glucocorticoid receptor gene, *Nr3c1*). We manipulated the natal brood size of zebra finches and sampled individuals in adulthood to explore the long-term impact of early-life conditions on the adult epigenetic landscape in a genome-wide, single-base resolution. We identified 149 DMSs with differential levels of DNAm between adult zebra finches raised in small and large broods. The finding that epigenetic signatures affected by early-life conditions remain constant throughout adulthood is in line with the above-mentioned correlational findings in mammals, and collectively demonstrate that early-life conditions can have long-lasting effects on epigenetic patterns observable in adulthood across various species. The effects of DNAm are highly contextual, i.e. the suppressing effects of DNAm of the promoter region is well established in multiple species including the zebra finch (Steyaert et al., 2016). However, only a single DMS was identified was in the promoter region of the *STX11* gene, which is known to play an important role in immune system function (Rudd et al., 2006) and pathogenesis of Peripheral T cell lymphomas (Yoshida et al., 2015). The elevated degree of DNAm of individuals raised large broods in the promoter region of *STX11* suggests suppressed gene expression that can lead to impaired immune function and increased risk of tumorigenesis. Other DMSs were spread across the genome, with the majority located in introns with a proportion significantly higher than expected by chance. Although the functional consequences of intron DNAm are less well known, research in fishes has demonstrated an inverse relationship between DNAm of the first intron and gene expression (Anastasiadi et al., 2018), pointing to a potential mechanism through which differential DNAm of intronic regions may also affect gene expression. Moreover, DNAm in introns can affect alternative splicing patterns (Lev Maor et al., 2015), leading to the production of different protein isoforms with diverse biological functions. We identified DMSs in four introns in the *SMYD3* gene, whose expression has been found to lead to poor prognosis in various types of cancer (Giakountis et al., 2017). Our results suggest a potential importance of DNAm in introns in the phenotypic effects of early-life stress in our population.

The identified enriched GO terms associated with the genes in or near which DMS we found were related to cell growth, division, and differentiation, regulation of inflammatory response, muscle contraction and neuronal function. The potential functional implications of the observed differential methylation on adult fitness prospects are discussed below.

One of the enriched GO terms identified is associated with the regulation of inflammatory response. The DNAm of genes involved in regulating inflammatory responses can act as an epigenetic regulator of the frequency and intensity of the innate and adaptive immune responses to disease, pathogens or injury (Surace & Hedrich, 2019). Parasite and disease resistance are linked to sexual selection and fitness in birds (Hamilton & Zuk, 1982; Westerdahl et al., 2012), highlighting how differences in gene expression in these areas mediated by DNAm can result in phenotypic variation resulting from early-life stress. Previous research did not find brood size effects on adult innate immune function of adult zebra finches (Driessen et al., 2021), but due to the complexity of the immune system this does not exclude the possibility that other aspects of the immune system are shaped by early-life stress.

Three of the identified significantly enriched GO terms were associated with muscle contraction, which is important for locomotion including flight (Biewener, 2011). Muscle contraction of the respiratory-vocal system aids birds in respiration and singing (Schmidt & Wild, 2014), which are vital for survival and reproduction respectively. The *CHRM3* gene, which is part of six significantly enriched GO terms, is involved in song learning in the zebra finch (Asogwa et al., 2018), and, in line with this finding natal brood size was demonstrated to affect song accuracy (Holveck et al., 2008). Another identified enriched GO term was associated with Ras GTPase signaling, which regulates multiple biological processes by acting as molecular switch near the plasma (Bourne et al., 1990;Ahearn et al., 2012). Moreover, Ras signal transduction also plays an important role in aging (Borrás et al., 2011; Slack et al., 2015) and differential DNAm in or near genes involved in Ras GTPase signaling, influenced by brood size during early development, could alter aging trajectories and contribute to the observed shorter lifespans of individuals reared in large broods (Briga et al., 2017). Sepers et al. (2021) also reported a DMS in the promoter of a GTPase activity regulating gene between nestling great tits in reduced and enlarged broods, suggesting that early-life stress can broadly affect GTPase signaling in passerines.

Jimeno et al. (2019) reported long-term effects of natal brood size on DNAm in or near the glucocorticoid receptor gene (*Nr3c1*) in zebra finches, with cascading effects on receptor expression and glucocorticoids. Methylation differences at *Nr3c1* CpG sites were less than 10% (see Figure 2 in Jimeno et al., 2019), and thus well below the 25% DMS threshold, and indeed we did not identify these sites as DMSs. However, did detect four CpG sites shared between the two data sets, all with elevated DNAm in individuals raised in large broods. The DNAm differences were also similar in magnitude, closely mirroring the patterns reported by Jimeno et al. (2019). The reproducibility of this pattern strengthens support for the hypothesis that methylation of the *Nr3c1* gene regulatory region underlies long-term phenotypic effects of natal brood size. This finding also highlights the importance of subtle epigenetic changes, given their apparent consistency and biological significance.

**Figure 2.**
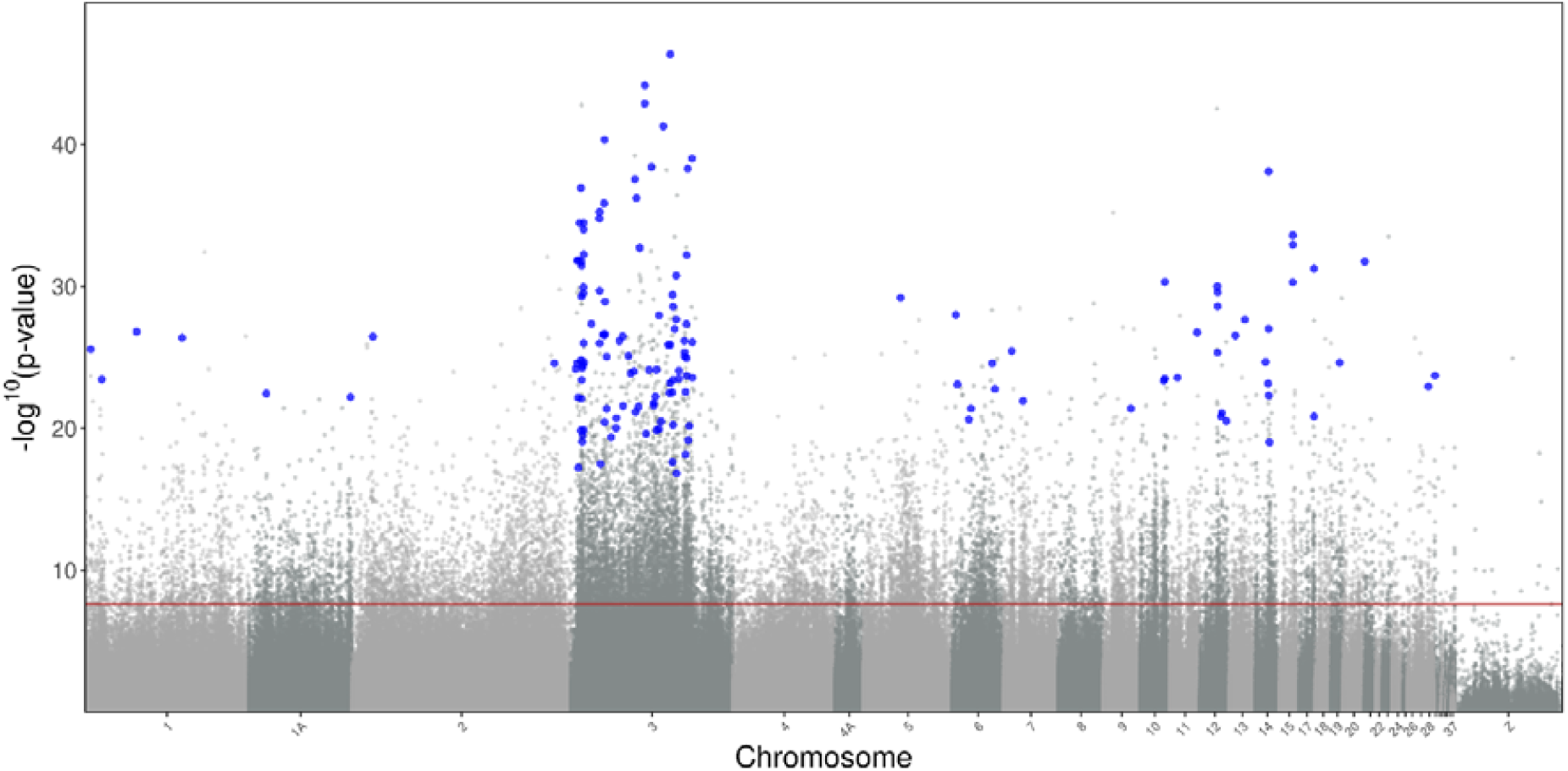
Manhattan plot of Bonferroni-corrected p-values across the genome. Each point represents a CpG site tested, with light and dark grey indicating non-significant differences and blue indicating biologically significant differences in DNAm between individuals raised in small and large broods. The red horizontal line marks the genome wide Bonferroni-corrected significance threshold of -log10(0.05/2,052,610)=7.61. Light and dark grey are used interchangeably to distinguish between the chromosomes and CpG sites are represented in their actual location within each chromosome.

**Figure 3.**
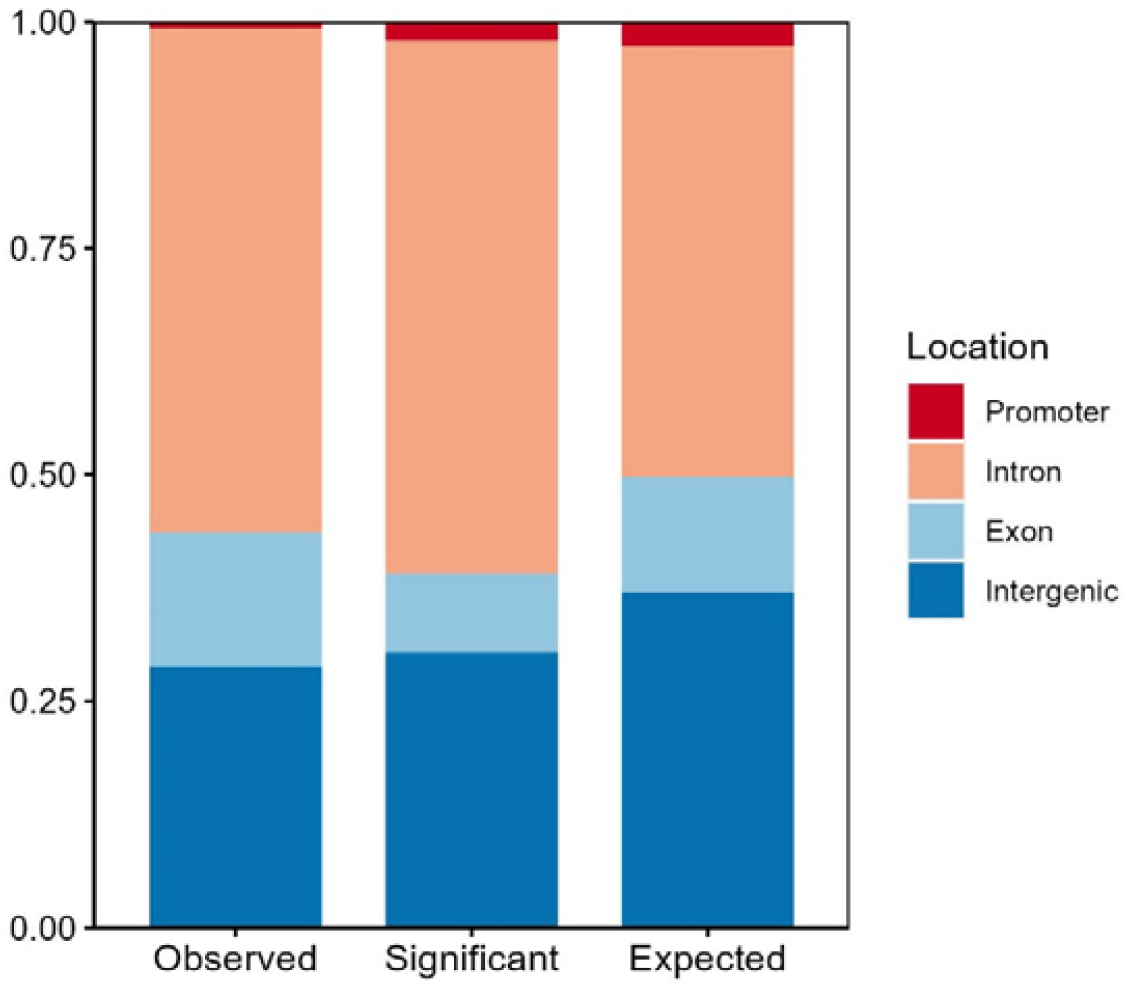
Distribution of three selections of CpG sites over different annotation categories. Observed: the 149 identified DMS, Significant: all CpG sites where DNAm differed significantly between individuals raised in small or large broods (Bonferroni-corrected, n=16,529), Expected: proportion of all CpG sites in each annotation category that was retained after filtering (n=2,052,610). DMS were significantly more often than expected by chance in introns, and less often than expected by chance in intergenic regions.

To our best knowledge, Sepers et al. (2021) conducted the only other study that explored CpG-site-specific differences in DNAm in relation to manipulated natal brood size. In our case we studied adults, whereas their study was restricted to the nestling stage. Sepers et al. (2021) identified 32 DMSs between great tit nestlings in reduced and enlarged broods, considerably fewer than in our study (149). The most immediate explanation for the difference between the two studies is likely because Sepers et al. (2021) employed reduced representation bisulfite sequencing and thus only covered a very small fraction of the overall genome. There was no overlap in genes containing DMSs detected by Sepers et al. (2021) and this study. This lack of overlap might be due to early-life stress effects on DNAm being life-stage (Wilks Siller et al., 2024) or species-specific. Studies covering both life stages are required to resolve this issue.

Our findings provide the foundation for several future research directions. The effects of brood size on phenotypic traits in some cases have been found to be sex-dependent in zebra finches (e.g. mortality, de Kogel, 1997; metabolic rate, Verhulst et al., 2006) and to expand on the present findings, investigation of sex-specific epigenetic responses to a manipulated brood size is of interest. Moreover, while the effects of DNAm in promoter regions on gene expression are well-established, the functional consequences of DNAm outside of the promoter regions remain poorly understood. Therefore, there is a need of experimental studies assessing how individual DMS influence gene expression, taking into account their specific genomic locations. Additionally, to assess the biological significance of DMS, there is a need to identify links between DMS and phenotypic traits known to differ between individuals raised in different brood sizes such as growth (Tangili et al., 2025) and mortality (Briga et al., 2017; de Kogel, 1997).

In summary, to our knowledge, we provided the first comprehensive characterization of how early-life conditions affect adult DNAm at a single-base resolution in an avian species. Our analysis revealed DMS between the two treatment groups located in or near genes whose differential DNAm patterns we hypothesize contribute to phenotypic variation and, consequently, the previously observed differential fitness outcomes. Our findings also suggest that, similarly to mammals, DNAm can mediate the long-term effects of early-life conditions on adult fitness of avian species, consistently shaping phenotypic quality during an individual’s life.

## Acknowledgements

We thank the Center for Information Technology of the University of Groningen for their support and for providing access to the Hábrók high performance computing cluster. We express our gratitude to Ellis Mulder for her help in the laboratory. We also thank the animal caretakers of the University of Groningen as well as numerous students whose invaluable help made this project possible. MB thanks the funding from the Turku Collegium for Science, Medicine and Technology.

## Ethics approval

All methods and experiments detailed in this manuscript were performed under the approval of the Central Committee for Animal Experiments (Centrale Commissie Dierproeven) of the Netherlands, under licenses AVD1050020174344.

## Data Accessibility and Benefit-Sharing

Raw reads of the whole methylomes of all samples are available at the National Center for Biotechnology Information under BioProject ID PRJNA1108628. The bioinformatics code is available in https://datadryad.org/dataset/doi:10.5061/dryad.wm37pvmw8, the R code and intermediate data files used in this study are available in https://github.com/tangilim/ZF_DNAm_BroodSize_DMS. The R code and intermediate data files can also be made available on Dryad after submission.

## Author contributions

SV designed the study. MB and BJ performed the brood size manipulations and collected the blood samples. MT performed the bioinformatic analysis, DMSs identification and statistical analysis. MT and SV wrote the manuscript with input from all authors. All authors approved the final version of the manuscript.

