## Supplementary Information for "Long-term effects of early-life adversity on DNA methylation in zebra finches"

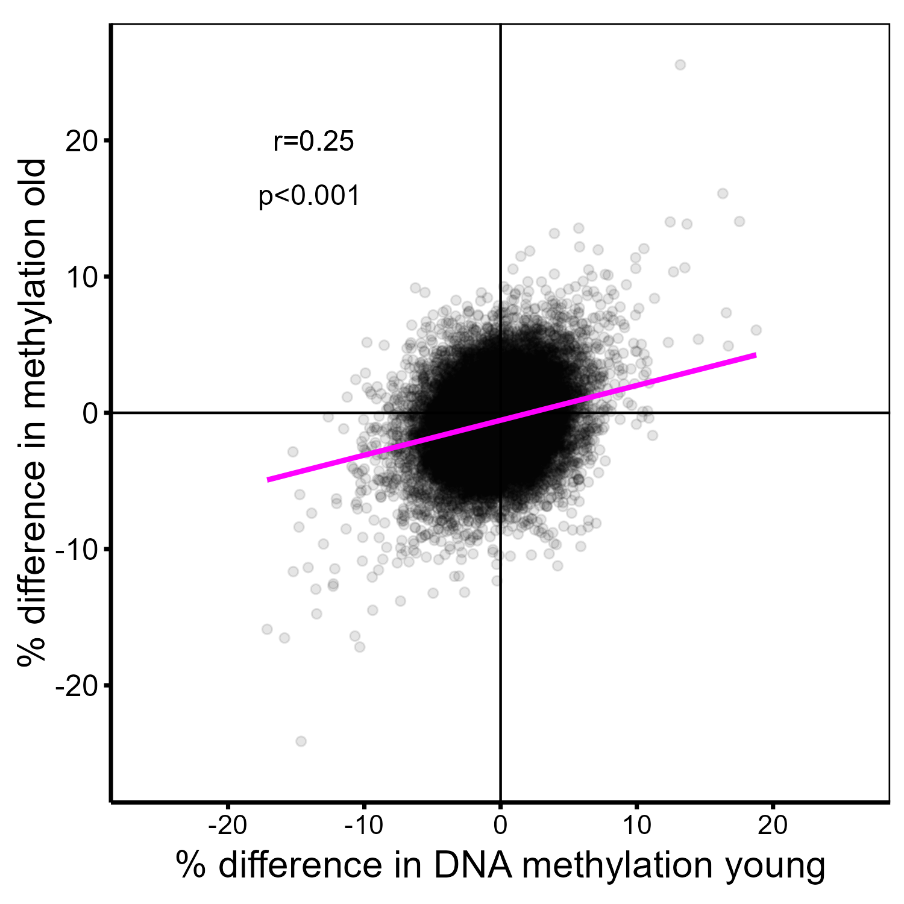

*FigS1. Percent DNA methylation difference between individuals raised in small vs. large broods in old against in young individuals. Each data point represents a CpG site.*

*Table S1. Exact position, methylation difference and annotation information for all DMS identified between individuals raised in small and large broods. A positive methylation difference value denotes higher methylation in large broods while a negative methylation difference value denotes higher methylation in small broods.*

| **Chromosome** | **Position** | **DNAm difference (large broods- small broods)** | **p-value** | **Region** | **Gene name** |
| --- | --- | --- | --- | --- | --- |
| 1 | 3998144 | -28.65 | 2.66E-26 | Intron | MAP3K7CL |
| 1 | 11643251 | 26.81 | 3.53E-24 | Intron | LOC115493364 |
| 1 | 35913092 | 27.65 | 1.53E-27 | Intergenic | SMCO4 |
| 1 | 67516484 | -25.84 | 4.09E-27 | Exon | SACS |
| 1A | 11871457 | -25.61 | 3.48E-23 | Intergenic | PTPN12 |
| 1A | 70280148 | 25.67 | 6.14E-23 | Intron | DDX11 |
| 2 | 14232709 | -27.83 | 3.39E-27 | Intergenic | NRP1 |
| 2 | 140208530 | -27.67 | 2.62E-25 | Intergenic | LOC116807906 |
| 3 | 3343580 | 29.15 | 6.39E-25 | Intron | BLK |
| 3 | 4201791 | -27.09 | 2.57E-25 | Intergenic | TFAP2B |
| 3 | 4347100 | -25.81 | 1.52E-32 | Intergenic | TFAP2B |
| 3 | 4997500 | -27.28 | 6.81E-23 | Exon | LOC105759121 |
| 3 | 5171398 | -25.34 | 5.91E-18 | Intergenic | LOC121469652 |
| 3 | 5348540 | -25.77 | 1.47E-32 | Intron | SUPT3H |
| 3 | 6060941 | 34.40 | 3.27E-35 | Exon | MEP1A |
| 3 | 7018989 | 32.27 | 1.15E-37 | Intergenic | EVA1A |
| 3 | 7045567 | -26.00 | 1.38E-20 | Intergenic | MRPL19 |
| 3 | 7045568 | -29.07 | 1.63E-25 | Intergenic | MRPL19 |
| 3 | 7243045 | -26.87 | 1.76E-32 | Intron | FKBP1B |
| 3 | 7281045 | -25.71 | 7.94E-23 | Intergenic | LOC100223718 |
| 3 | 7439076 | -29.01 | 5.20E-30 | Intergenic | ATAD2B |
| 3 | 7498888 | -30.17 | 3.48E-32 | Intron | ATAD2B |
| 3 | 7555610 | -25.08 | 3.93E-24 | Intron | LOC121469571 |
| 3 | 7905060 | 25.95 | 8.59E-20 | Intergenic | KLHL29 |
| 3 | 7937801 | -27.43 | 5.42E-25 | Intergenic | KLHL29 |
| 3 | 8185836 | -25.87 | 2.87E-20 | Intergenic | KLHL29 |
| 3 | 8466804 | 27.53 | 3.57E-25 | Intergenic | LOC115494710 |
| 3 | 8601210 | -26.04 | 3.13E-30 | Intergenic | LOC115494710 |
| 3 | 8652428 | -26.23 | 1.15E-30 | Intergenic | LOC115494710 |
| 3 | 8812202 | -29.35 | 3.24E-35 | Intergenic | LOC115494710 |
| 3 | 8812365 | -25.65 | 1.30E-20 | Intergenic | LOC115494710 |
| 3 | 8812366 | -29.63 | 9.84E-35 | Intergenic | LOC115494710 |
| 3 | 8835705 | 25.99 | 9.81E-27 | Intergenic | LOC115494710 |
| 3 | 8968189 | 27.30 | 5.51E-33 | Intergenic | LOC115494710 |
| 3 | 9079661 | -26.15 | 2.11E-25 | Intergenic | TDRD15 |
| 3 | 14189631 | -25.29 | 4.30E-28 | Intron | ROCK2 |
| 3 | 19831862 | 33.72 | 1.63E-35 | Exon | FAM110C |
| 3 | 19932912 | -30.24 | 5.62E-36 | Intron | FBXO25 |
| 3 | 19934605 | -25.98 | 9.93E-27 | Intron | FBXO25 |
| 3 | 19947392 | 27.18 | 2.04E-30 | Intron | FBXO25 |
| 3 | 20561680 | 25.39 | 3.13E-18 | Intron | CLN8 |
| 3 | 22270522 | 27.55 | 2.38E-27 | Intron | LOC115494608 |
| 3 | 23038815 | -32.56 | 1.39E-36 | Intron | MCPH1 |
| 3 | 23040039 | -26.46 | 2.78E-27 | Intron | MCPH1 |
| 3 | 23355826 | -30.31 | 4.65E-41 | Intron | CILK1 |
| 3 | 23429196 | 26.04 | 3.74E-21 | Intron | GCM1 |
| 3 | 23660719 | 27.68 | 2.34E-27 | Intron | LOC115494251 |
| 3 | 23670589 | 28.14 | 1.17E-29 | Intron | LOC115494251 |
| 3 | 24777783 | 28.05 | 9.06E-26 | Intron | DST |
| 3 | 24780620 | 25.87 | 4.18E-22 | Intron | DST |
| 3 | 27909952 | 25.11 | 4.36E-20 | Intergenic | LOC121469676 |
| 3 | 31315730 | 26.73 | 9.48E-21 | Intergenic | IMPG1 |
| 3 | 31485634 | 25.21 | 1.87E-21 | Intergenic | HTR1B |
| 3 | 33678489 | 25.16 | 6.99E-27 | Intron | DOP1A |
| 3 | 35956494 | 27.47 | 3.22E-27 | Exon | LOC100230156 |
| 3 | 36205343 | -26.12 | 2.68E-22 | Intron | CASP8AP2 |
| 3 | 39975845 | -25.16 | 7.90E-26 | Intron | FBXL4 |
| 3 | 41457260 | 25.30 | 1.34E-24 | Intron | GRIK2 |
| 3 | 43787719 | -27.40 | 9.55E-25 | Intron | LOC115494303 |
| 3 | 44396974 | -31.06 | 2.74E-38 | Intron | LOC115494307 |
| 3 | 44812966 | -26.77 | 6.96E-22 | Intron | ZBTB24 |
| 3 | 45550178 | 29.00 | 6.15E-37 | Exon | MFSD4B |
| 3 | 46926456 | -25.68 | 3.02E-22 | Intergenic | LOC115494314 |
| 3 | 47744976 | -29.97 | 1.91E-33 | Intron | NT5DC1 |
| 3 | 51298310 | 31.86 | 6.95E-45 | Intron | LOC115494610 |
| 3 | 51298333 | 27.78 | 1.33E-43 | Intron | LOC115494610 |
| 3 | 52135458 | -26.48 | 2.35E-20 | Intergenic | CENPW |
| 3 | 54223239 | -29.30 | 7.54E-25 | Intron | MSH5 |
| 3 | 55878810 | 26.21 | 3.89E-39 | Intergenic | HBS1L |
| 3 | 57408483 | 25.80 | 2.40E-22 | Intron | ESR1 |
| 3 | 57593145 | 26.91 | 1.63E-22 | Exon | SYNE1 |
| 3 | 58746360 | 26.57 | 5.59E-23 | Exon | SCAF8 |
| 3 | 59411877 | -27.97 | 1.34E-20 | Intergenic | LOC121469719 |
| 3 | 59411878 | -26.32 | 7.26E-25 | Intergenic | LOC121469719 |
| 3 | 61127309 | 26.40 | 1.10E-20 | Intron | HIVEP2 |
| 3 | 61127310 | 26.32 | 1.08E-28 | Intron | HIVEP2 |
| 3 | 62579221 | -27.13 | 3.32E-21 | Intron | REPS1 |
| 3 | 64108244 | 25.26 | 5.20E-42 | Promoter | STX11 |
| 3 | 67556957 | -25.53 | 1.29E-26 | Intergenic | CHRM3 |
| 3 | 68325858 | 27.63 | 3.14E-23 | Intron | MTR |
| 3 | 68825022 | 25.09 | 6.39E-24 | Exon | GGPS1 |
| 3 | 68942377 | -29.96 | 1.28E-26 | Intergenic | LOC115494755 |
| 3 | 68942378 | -34.33 | 4.39E-47 | Intergenic | LOC115494755 |
| 3 | 70390894 | 25.51 | 2.85E-23 | Intron | DISC1 |
| 3 | 70507016 | -25.98 | 2.39E-18 | Intergenic | TSNAX |
| 3 | 70564768 | 29.10 | 3.84E-30 | Exon | EXOC8 |
| 3 | 70815143 | 26.44 | 4.01E-24 | Exon | LOC101233512 |
| 3 | 70881600 | 28.88 | 2.60E-29 | Intron | NUP133 |
| 3 | 70881601 | 26.46 | 5.77E-21 | Intron | NUP133 |
| 3 | 72080988 | 25.62 | 9.64E-28 | Intron | WDR27 |
| 3 | 73093190 | -29.68 | 2.09E-28 | Intron | AFDN |
| 3 | 73093191 | -29.89 | 1.67E-31 | Intron | AFDN |
| 3 | 73293232 | 27.04 | 1.47E-17 | Intergenic | UNC93A |
| 3 | 74946263 | -26.58 | 8.61E-25 | Intergenic | LOC115494398 |
| 3 | 74957975 | -25.26 | 3.17E-24 | Intergenic | LOC115494398 |
| 3 | 78656871 | -25.01 | 6.93E-27 | Intron | RMDN2 |
| 3 | 79019801 | 25.38 | 8.68E-26 | Intron | LOC105759092 |
| 3 | 79047173 | 26.83 | 4.27E-26 | Intron | LOC105759092 |
| 3 | 79515663 | 25.75 | 6.89E-19 | Intron | LOC115494415 |
| 3 | 79570323 | -25.39 | 2.69E-23 | Intron | ZNF318 |
| 3 | 80141895 | -28.22 | 4.56E-28 | Intron | SMYD3 |
| 3 | 80181379 | 26.26 | 1.12E-25 | Intron | SMYD3 |
| 3 | 80235838 | -25.11 | 1.92E-24 | Intron | SMYD3 |
| 3 | 80282163 | -26.91 | 6.15E-33 | Intron | SMYD3 |
| 3 | 81053803 | -31.49 | 4.95E-39 | Intergenic | ZBTB18 |
| 3 | 81429092 | -25.25 | 6.99E-20 | Exon | CEP170 |
| 3 | 82216823 | 26.60 | 6.67E-21 | Intergenic | LOC115494427 |
| 3 | 84068277 | 28.46 | 9.88E-40 | Intergenic | RIN2 |
| 3 | 84285545 | 29.19 | 8.43E-27 | Intron | RALGAPA2 |
| 3 | 84285546 | 25.15 | 2.76E-24 | Intron | RALGAPA2 |
| 5 | 25768618 | 28.79 | 6.17E-30 | Intron | CAPN3 |
| 6 | 2344488 | -27.59 | 9.89E-29 | Intron | LOC100220618 |
| 6 | 3527947 | 27.31 | 8.07E-24 | Exon | DNAJC9 |
| 6 | 11315632 | 27.04 | 2.42E-21 | Intron | LOC115495760 |
| 6 | 12896255 | 26.49 | 4.10E-22 | Intron | DLG5 |
| 6 | 27609082 | 25.92 | 2.62E-25 | Intron | CCDC186 |
| 6 | 29505768 | -26.09 | 1.67E-23 | Intron | RAB11FIP2 |
| 7 | 5549871 | -26.41 | 3.63E-26 | Intron | SLC49A4 |
| 7 | 13366643 | -26.93 | 1.16E-22 | Exon | B3GALT1 |
| 9 | 19019209 | -25.98 | 4.10E-22 | Exon | KCNMB2 |
| 10 | 16354714 | -27.54 | 4.45E-24 | Intron | LOC115496633 |
| 10 | 17000251 | 25.50 | 3.10E-24 | Exon | LOC115496570 |
| 10 | 17060482 | 31.46 | 4.83E-31 | Intron | LRRC28 |
| 11 | 5317152 | 25.39 | 2.71E-24 | Intron | CTCF |
| 11 | 18959616 | -26.17 | 1.72E-27 | Intron | KCTD15 |
| 12 | 11809741 | -29.10 | 9.96E-31 | Intron | DNASE1L3 |
| 12 | 11923636 | 27.58 | 4.60E-26 | Intron | LOC100218261 |
| 12 | 12050493 | 25.88 | 2.49E-30 | Intron | DNAH12 |
| 12 | 12050494 | 26.20 | 2.48E-29 | Intron | DNAH12 |
| 12 | 14366311 | -27.21 | 1.44E-21 | Intron | LOC115496928 |
| 12 | 14936178 | -25.81 | 8.07E-22 | Exon | GLT8D1 |
| 12 | 18085885 | -25.31 | 3.02E-21 | Intron | MIR2971 |
| 13 | 4017424 | 25.10 | 2.88E-27 | Intergenic | LOC115497102 |
| 13 | 10638530 | -25.57 | 2.20E-28 | Intron | MAPK9 |
| 14 | 6813807 | 25.88 | 2.14E-25 | Intron | FAHD1 |
| 14 | 8731302 | 26.34 | 6.98E-24 | Intron | LOC100227445 |
| 14 | 9013940 | 30.03 | 7.73E-39 | Intron | SNX29 |
| 14 | 9028062 | 26.48 | 9.73E-28 | Intron | SNX29 |
| 14 | 9061150 | -25.04 | 4.79E-23 | Intron | SNX29 |
| 14 | 9645048 | -25.13 | 9.63E-20 | Intron | MRTFB |
| 15 | 9499190 | -29.63 | 1.15E-33 | Exon | KIAA1671 |
| 15 | 9499192 | -29.71 | 5.08E-31 | Exon | KIAA1671 |
| 15 | 9499195 | -30.68 | 2.41E-34 | Exon | KIAA1671 |
| 17 | 10465455 | -27.17 | 1.47E-21 | Exon | LOC116809029 |
| 17 | 10465456 | -28.68 | 5.47E-32 | Exon | LOC116809029 |
| 19 | 5790153 | -27.30 | 2.41E-25 | Intron | RASL10B |
| 21 | 113753 | 27.46 | 1.80E-32 | Intron | C21H1orf159 |
| 28 | 2296515 | 26.23 | 1.13E-23 | Intron | EPS15L1 |
| 29 | 950179 | -26.71 | 1.88E-24 | Intron | SARNP |

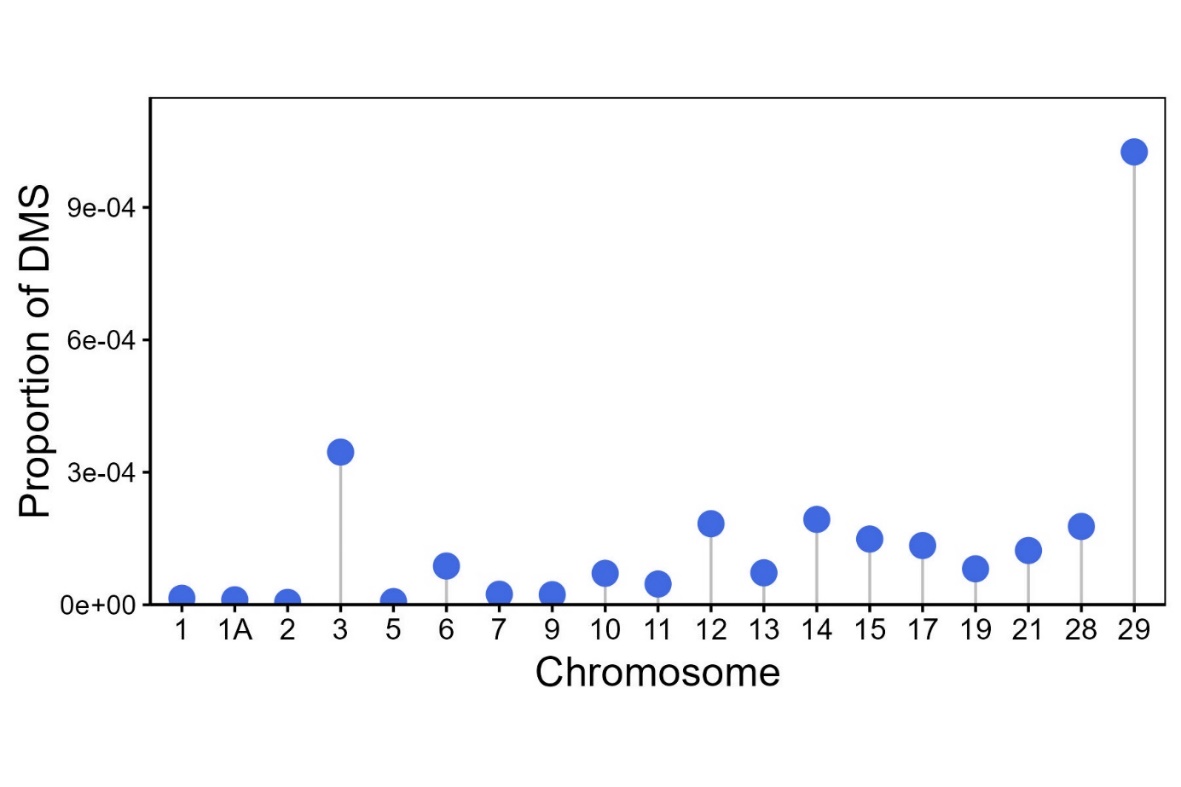

*Figure S2. Proportion of DMS sites out of all CpG sites retained after filtering for each chromosome in which a DMS was found.*

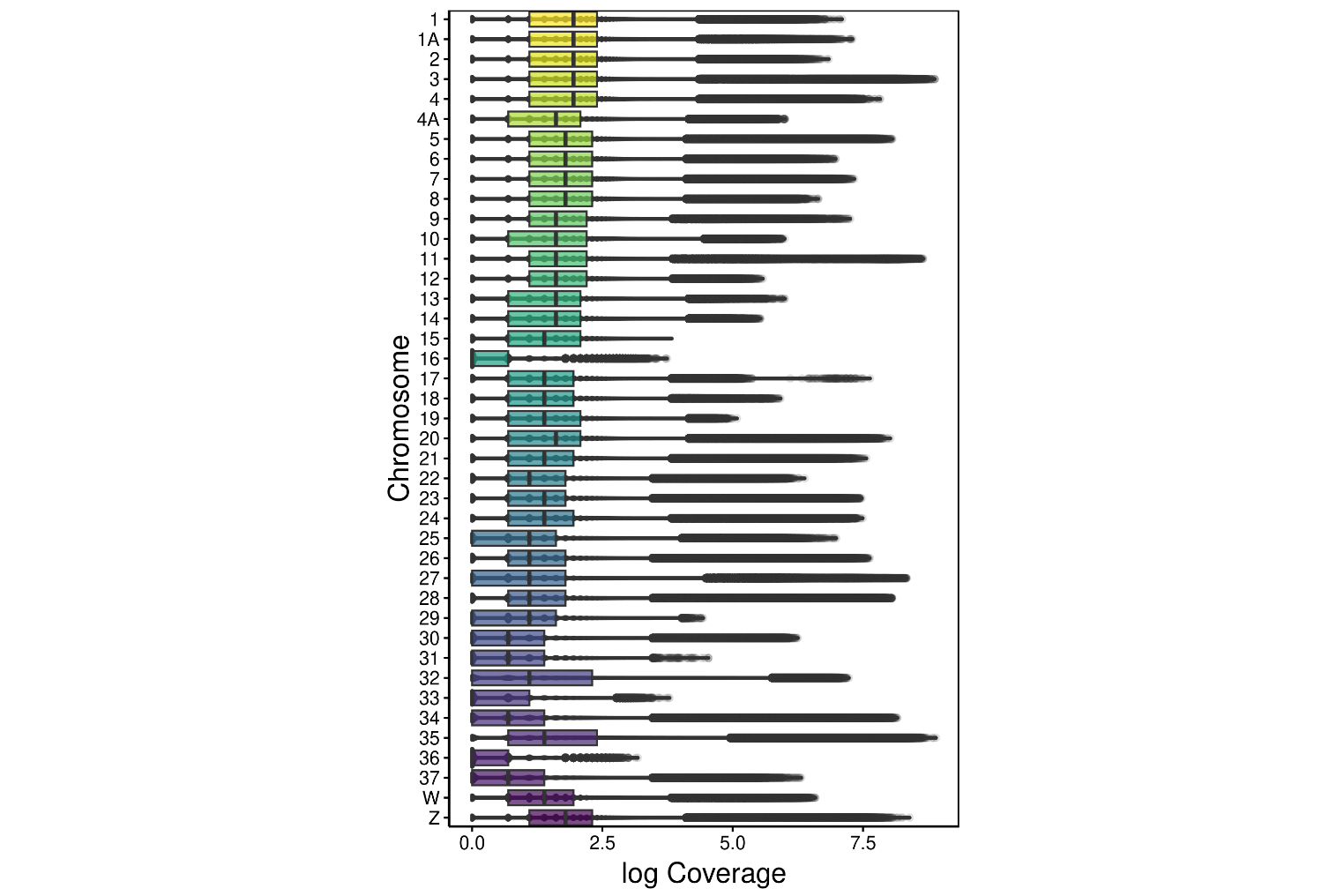

*Figure S3. Coverage (in log_10_ scale) per chromosome.*

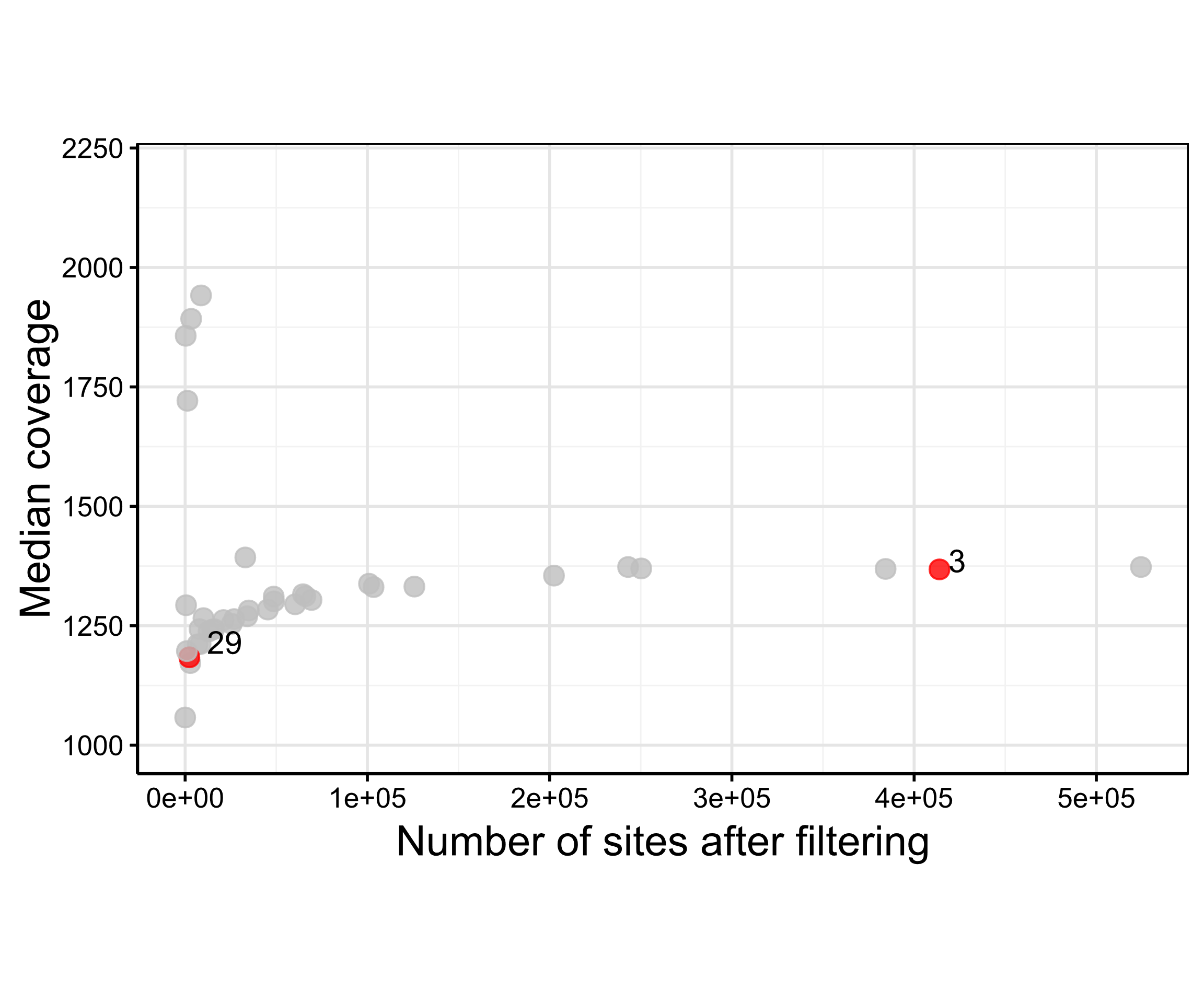

*Figure S4. Median coverage against the number of sites in each chromosome after filtering.*

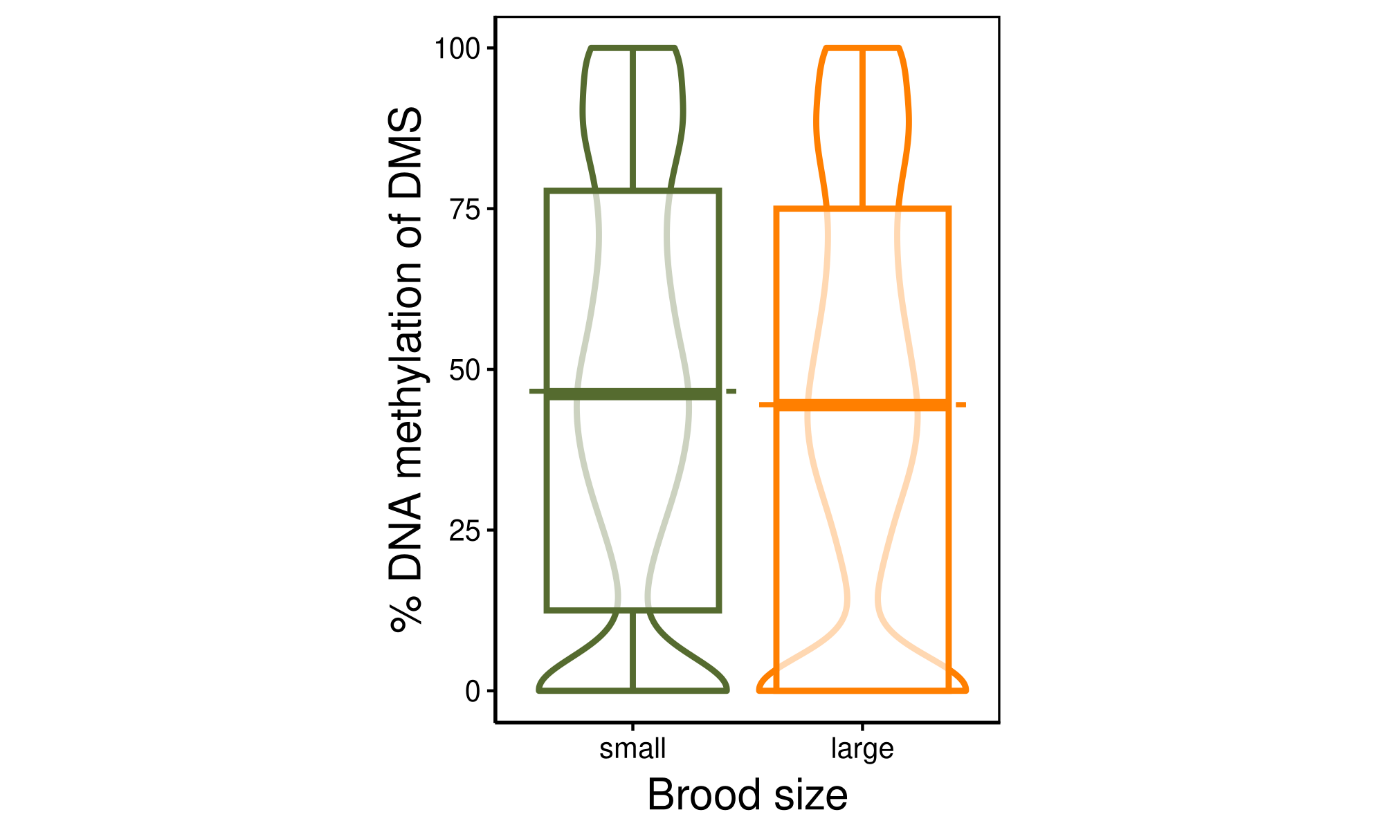

*Figure S5. Percentage of DNA methylation at identified DMS in small and large broods. Violin plot widths represent the data density, boxplots show the median and quartiles while dashed lines show the average percent DNA methylation level per brood size.*

Table S2. Enriched GO terms, descriptions, p-values and associated genes.

| **GO term ID** | **Description** | **FDR-corrected p-value** | **Associated Genes Found** | **Ontology** |
| --- | --- | --- | --- | --- |
| GO:0017016 | Ras GTPase binding | 0.01 | EXOC8, RAB11FIP2, ROCK2 | Molecular function |
| GO:0050727 | regulation of inflammatory response | 0.01 | DNASE1L3, ESR1, MCPH1 | Biological process |
| GO:0071103 | DNA conformation change | 0.01 | CTCF, DDX11, MCPH1 | Biological process |
| GO:0071824 | protein-DNA complex subunit organization | 0.02 | CENPW, CTCF, ESR1 | Biological process |
| GO:0003012 | muscle system process | 0.01 | CHRM3, FKBP1B, ROCK2 | Biological process |
| GO:0006936 | muscle contraction | 0.01 | CHRM3, FKBP1B, ROCK2 | Biological process |
| GO:0006939 | smooth muscle contraction | 0.00 | CHRM3, FKBP1B, ROCK2 | Biological process |
| GO:0030594 | neurotransmitter receptor activity | 0.01 | CHRM3, GRIK2, HTR1B | Molecular function |
| GO:0035690 | cellular response to drug | 0.01 | CHRM3, DDX11, HTR1B | Biological process |
| GO:0097060 | synaptic membrane | 0.02 | CHRM3, GRIK2, HTR1B | Cellular component |
